# Brainwide mesoscale functional networks revealed by focal infrared neural stimulation of the amygdala

**DOI:** 10.1101/2024.02.14.580397

**Authors:** An Ping, Jianbao Wang, Miguel Ángel García-Cabezas, Lihui Li, Jianmin Zhang, Katalin M. Gothard, Junming Zhu, Anna Wang Roe

## Abstract

The primate amygdala serves to evaluate emotional content of sensory inputs and modulate emotional and social behaviors; it modulates cognitive, multisensory and autonomic circuits predominantly via the basal (BA), lateral (LA), and central (CeA) nuclei, respectively. Based on recent electrophysiological evidence suggesting mesoscale (millimeters-scale) nature of intra-amygdala functional organization, we have investigated the connectivity of these nuclei using Infrared Neural Stimulation of single mesoscale sites coupled with mapping in ultrahigh field 7T functional Magnetic Resonance Imaging (INS-fMRI). Stimulation of multiple sites within amygdala of single individuals evoked ‘mesoscale functional connectivity maps’, allowing comparison of BA, LA and CeA connected brainwide networks. This revealed a mesoscale nature of connected sites, complementary spatial patterns of functional connectivity, and topographic relationships of nucleus-specific connections. Our data reveal a functional architecture of systematically organized brainwide networks mediating sensory, cognitive, and autonomic influences from the amygdala.

## Introduction

The primate amygdala evaluates the emotional salience of inputs from all sensory modalities and contributes to the elaboration of emotional and social behaviors^1,2^. Anatomical, electrophysiological, and behavioral studies indicate that the integration and dissemination of neural information across sensory-motor and decision networks is achieved by multiple functional processing loops connecting the amygdala to a wide array of cortical and subcortical targets^3–5^. However, while much anatomical data exists on amygdala’s direct connections with other brain areas, these data do not reveal the functional *brainwide* networks within which amygdala selectively mediates its complex array of sensory, motor, cognitive, and physiological influence. Functional resting state data provide more information on brainwide networks^6,7^, but generally do not address connections patterns at mesoscale (millimeter-scale). Investigation into functional networks at mesoscale is of interest given the unique mesoscale cortical architecture in human and nonhuman primates previously described as ‘columnar’ or ‘modular’, raising questions regarding how amygdala interfaces with such modular organizations.

Three subdivisions of the amygdala which have received significant anatomical study are the Basal (BA), Lateral (LA), and Central (CeA) nuclei^8–12^. CeA projects primarily to targets in the basal forebrain, hypothalamus, and brainstem^13^. BA projects to large swaths of the frontal, insular, temporal, and visual cortex, as well as claustrum and cingulate cortex^8,14^. LA gives rise to projections to orbitofrontal, cingulate, and insular cortex, as well as hippocampal areas^15^. Electrical stimulation of nuclear-specific targets in the basal vs lateral human amygdala combined with neuroimaging have also revealed multiple networks characterized by distinct spatiotemporal patterns of activation ^16^. Thus, functional distinctions in connectivity associate CeA, BA, and LA predominantly with autonomic, cognitive, and multisensory circuits, respectively^6,17^.

Whether there are further functional distinctions within each of the CeA, BA, and LA nuclei is less clear. Anatomical tracing studies have suggested the presence of topographic gradients within BA, connecting the magnocellular (dorsal) division of BA with occipital visual areas and more parvicellular (ventral) division with more anterior visual areas^14^. Evidence from mice and macaques also suggest heterogeneity of cell types^18^. Functionally, evaluation using local field potential and current source density recordings in Macaque amygdala has revealed a clear mesoscale (millimeters-scale) signature of intra-amygdala function and circuitry^19,20^, indicating possible presence of functional clustering. In fact, multi-site electrophysiological current source density recordings show that this diversity results in distinct (e.g., visual, tactile, auditory, and multisensory) mesoscale functional organization within the amygdala in behaving monkeys^19,20^, the integration of which is central to the interpretation of social facial communications^19^. This raises the exciting possibility that connectional relationships between the amygdala and brainwide networks are also organized at mesoscale.

A recent technological development has introduced a novel method for studying functional networks brainwide at mesoscale. This method, termed INS-fMRI, uses pulsed Infrared Neural Stimulation (INS) to stimulate submillimeter sites in the brain, leading to activation of connected sites which is mapped using high-resolution Functional Magnetic Resonance Imaging^21^. Unlike optogenetics, this method is non-viral and can be used to stimulate any submillimeter cluster of neurons in the brain via fine (200um diameter) optical fibers; functional activation of connected sites are mapped at the full brain scale by recording Blood Oxygen Level Dependent (BOLD) signals (for review of INS, see^22–24^; for membrane capacitance effects see^25^; for safety and damage thresholds in primates, see^26,27^. Importantly, as INS reveals functional connectivity extending up to two synapses from the stimulation site ^21^, it evokes a broader, yet highly specific, set of network activations, beyond what traditional anatomical tracing provides^28,29^. INS thus provides distinct benefits for high resolution circuit mapping^30,31^, something especially relevant for columnar organization in primate brains. Another important advantage of this approach is that multiple sites can be stimulated within a single animal, providing a rich dataset of multiple networks whose organization and mutual relationships can be compared.

Using this method, we previously established proof-of-principle that INS-fMRI in Macaque amygdala BA reveals statistically significant, mesoscale connectivity with connected sites in the cingulate, insular, and association sensory cortex^28,29^. Here, to study more systematically the connectivity of mesoscale sites in each of the BA, LA, and CeA nuclei, we have mapped the cortical connectivity of stimulation sites within individual Macaque monkeys, permitting comparison of within-monkey networks at brainwide scale. The resulting activation maps (1 site: 1 network) reveal that functional specificity is achieved via sets of mesoscale activations within single cortical areas, and that multi-modal cognitive, sensory, and autonomic influences are arranged in spatially organized patterns of topographic or interdigitating connectivity. These findings suggest that amygdala influence, previously proposed as heterogeneous processing loops, is embodied in an architecturally organized cortical mesoscale interface.

## Results

### Overview

The purpose of this study was to examine the organization of connections between the amygdala and various cortical areas in individual Macaque monkeys. We note that, compared with most anatomical studies where tracer injections can span several millimeters, our stimulation is significantly more focal, activating, with intensities used, a volume of tissue <1mm^3^ ^32^. A major strength of the INS method is that networks activated by the stimulation of multiple sites can be compared within an individual. A weakness is that, due to the focal nature of the stimulation, we are sampling a small volume of the total amygdala. This study presents a sampling of nuclei CeA, BA, and LA (**Fig. 2A**) and reveals the mesoscale aspect of their connection patterns (**Fig. 2B**). Our rationale for analysis of the data begins with matrix-based comparison of known anatomy and presence or absence of INS-evoked connections (**Fig. 3A**). Further, as functional connections include both ‘first synapse’ and ‘second synapse’ connections, the functional connectivity networks are expected to be greater than the anatomical connectivity networks. Note that, because INS-fMRI is biased in the ‘anterograde’ direction, these connections reflect more strongly amygdalofugal functional connectivity (**Fig. 3B&C**). We then examine, within each of the cortical areas (which span limbic, sensory, motor, cognitive, and prefrontal cortical areas), the spatial distribution of connections (**Fig. 3D-H**). The examples shown in **Fig. 4** highlight connections patterns in limbic cortical areas. **Fig. 5** and **Fig. 6** illustrate examples of topographic and interleaved connection patterns.

### INS of the amygdala reveals remote connections at mesoscale

By inserting a 200μm diameter optical fiber through a preinstalled grid into the amygdala (**Fig. 1C**), we stimulated discrete sites in the right amygdala of monkey K and determined the nuclear location of the stimulation site with a 0.3mm precision (see Methods and previous study^28^). Periodic trains of pulsed infrared neural stimulation at the stimulated site (**Fig. 1D**) evoked BOLD signal response (**Fig. 1E**). As previously shown^21,28,29^, functional connectivity at remote sites were evaluated by correlation to the stimulation site (see Methods). **Fig. 1F-I** presents an example of a connected site in the frontal lobe with a timecourse with statistically significant correlation with the INS stimulation (**Fig. 1J**). A correlation value was obtained for every voxel in the brain; only voxels of high statistical significance (T-test p<1×10^−3^, see Methods) were further studied. Reproducibility was evaluated using comparisons of half-runs (e.g., even vs. odd runs, **Fig. S2** and previous study^28^). Reliability of activation was also evaluated by examining different thresholds; generally, with lower p-values, activation sizes increased, but activations remained in a similar location (**Fig. S3** and previous study^29^), supporting the reliability of the activation location.

**Fig. 1.**
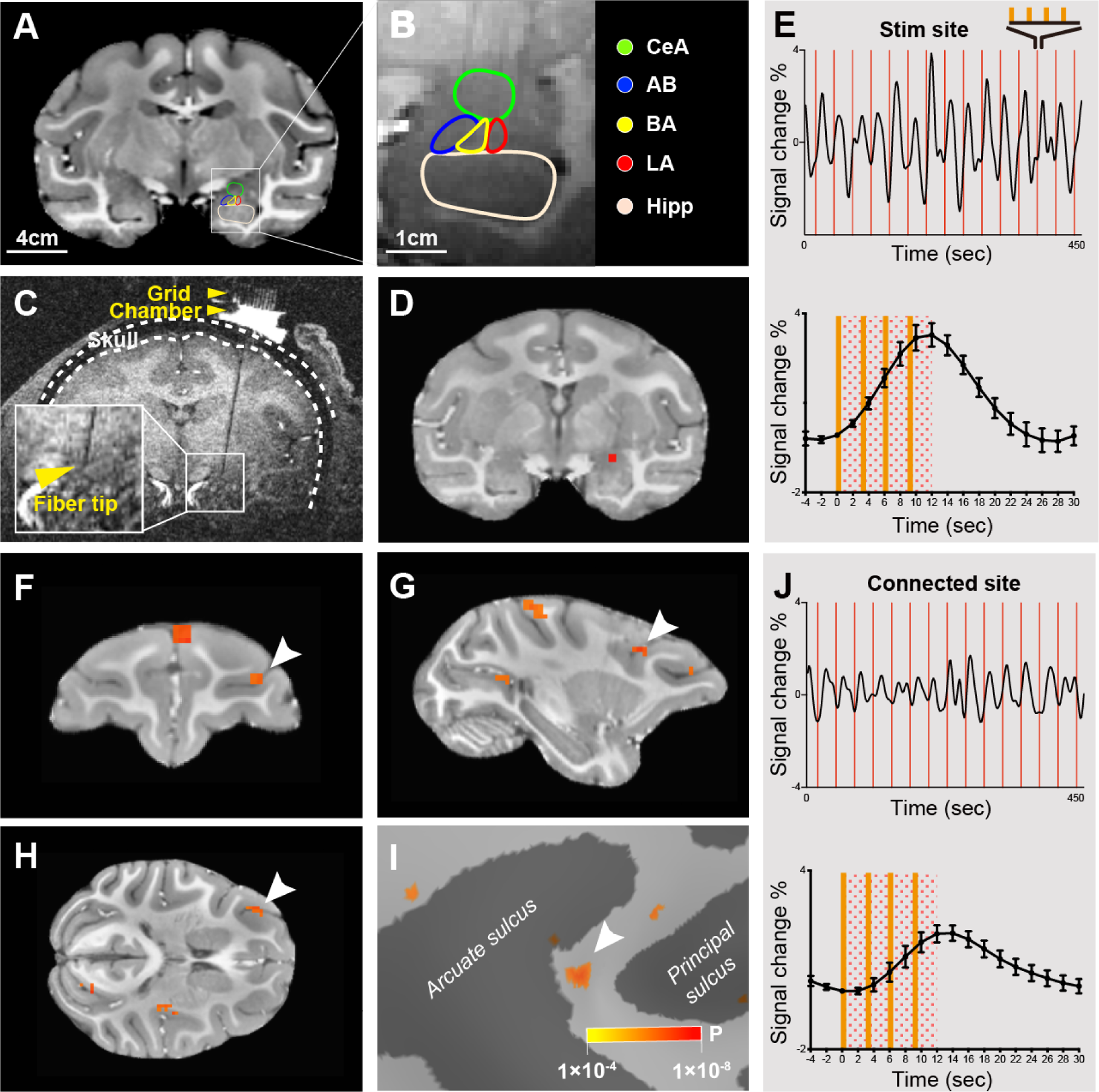
Identifying functionally connected sites in the brain following INS stimulation of single mesoscale sites the amygdala. (A) A coronal section through the caudal amygdala. (B) Parcellation of the amygdala at the most caudal site shown in A. CeA: central amygdala. AB: accessory basal amygdala. BA: basal amygdala. LA: lateral amygdala. Hipp: hippocampus. (C) Raw structural image indicating the optical fiber inserted through a grid in a chamber. (D) Activation at the laser tip in CeA (red voxel, intensity: 0.2 J/cm2, p<1×10-6). (E) BOLD time course at the laser tip in D. Above: 15 consecutive trials; Below: averaged time course (the dotted rectangle spans the duration of INS). Each red line: one trial of 4 pulse trains (see Methods). (F-H) Coronal, sagittal and horizontal view of a remote cluster activated in response to stimulation in D (p<1×10-4). (I) Activation cluster (white arrow) in F-H shown on inflated brain surface. (J) BOLD time course at connected cluster (white arrow) in F-H. Above: all 15 trials; Below: averaged time course.

### Distribution of global cortical connections from CeA, BA and LA

To obtain a comparison of amygdala stimulated networks, for each animal (Monkey K, Monkey M), we examined data acquired within a single animal (**Fig. 2A**, example shown of Monkey K: 6 sites in CeA, 3 sites in BA and 3 sites in LA) (**Fig. 2A**). Note that, unlike anatomical tracer injections which tend to fill more of the amygdala (e.g., a subdivision), this study has sampled very focal (millimeter-sized) locations in different parts of the amygdala. Voxels with significant p-values were selected for subsequent statistical analysis (see Methods and previous study^28^). As shown in **Fig. 2B & 2C**, activations from single site stimulation appeared patchy and mesoscale in size. The distribution had a sparse appearance and spanned multiple brain areas. This was seen consistently across every stimulation site in both Monkey K and Monkey M (examples shown in **Fig. 2C**). Although some activations at a single threshold were large (>10mm^2^, <10%), patch sizes from stimulation in CeA, BA, and LA were predominantly (64% of patches) less than 3mm^2^ in size (**Fig. 2B**, for monkey M see **Fig. S1**). Note that, while the size of mesoscale domains is dependent on the threshold used, the locations of these activations remain stable and distinct (see **Fig. S3**). Thus, what we show in this study are the activations that have the strongest functional connections (highest correlations) with the stimulation site, providing a view of the ‘backbone’ of a functional network. Nodes within this functional network may be modulated in size and strength during natural behavior.

**Fig. 2.**
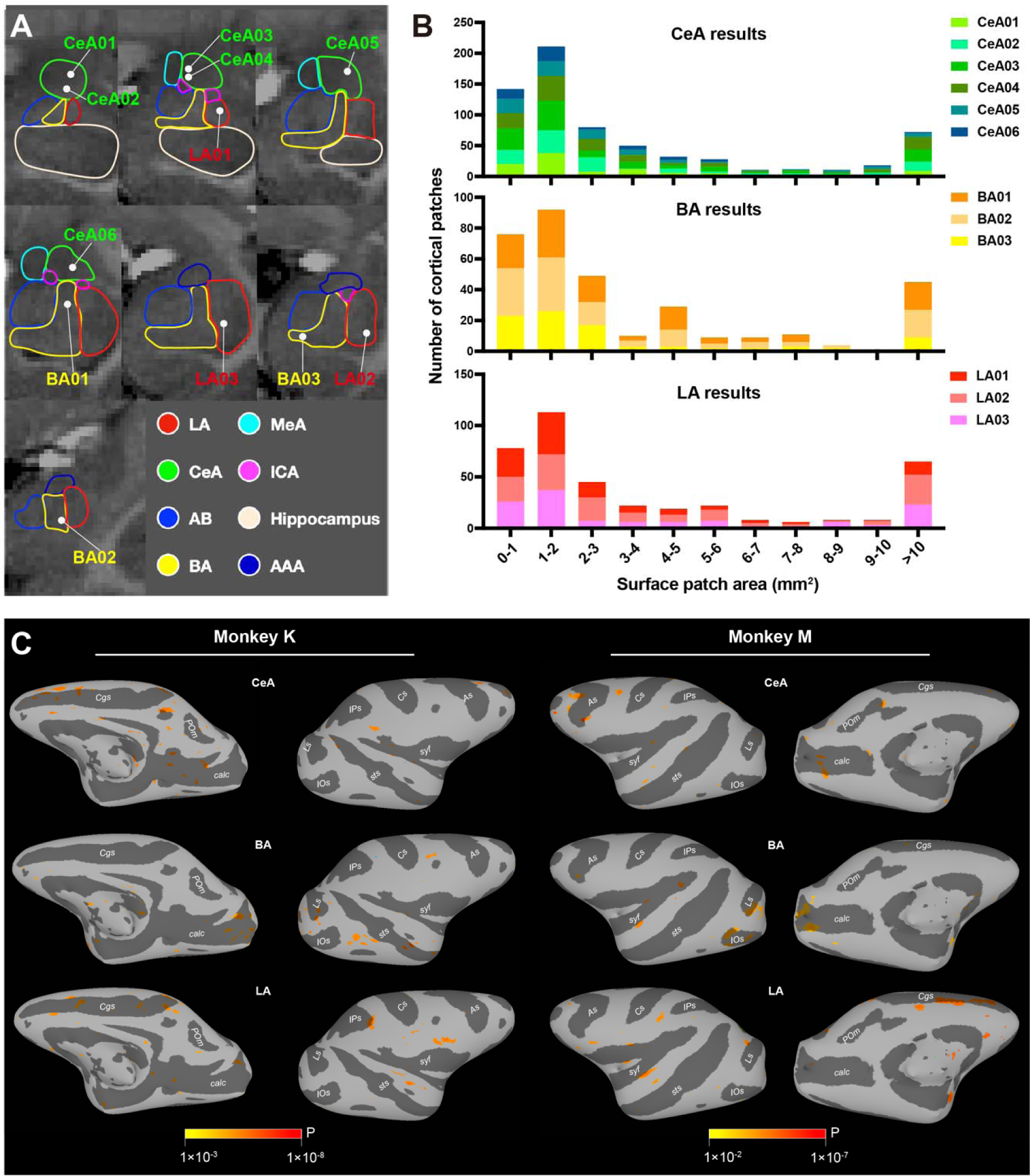
Mesoscale brainwide connections of the amygdala. (A) The white dots represent the INS stimulation sites in the right amygdala. CeA (green contour, 6 sites), BA (yellow contour, 3 sites), LA (red contour, 3 sites) LA: lateral amygdala. BA: basal amygdala. AB: accessory basal amygdala. CeA: central amygdala. MeA: medial amygdala. ICA: intercalated cell masses. AAA: anterior amygdala area. (B) The stacked histogram for patch size of brainwide cortical activations. The x axis represents the size of patches in millimeter square. The y axis represents the number of patches of different sizes. Each color represents a stimulation site in monkey K, namely 6 sites in CeA (upper row, shades of green), 3 sites in BA (middle row, shades of yellow) and 3 sites in LA (lower row, shades of red). (C) Whole brain activations evoked by single stimulation sites (1 site for each of CeA, BA and LA) mapped on inflated hemisphere (ipsilateral to the stimulation) of monkey K and monkey M. Both medial view and lateral view are presented. Ps: principal sulcus. As: arcuate sulcus. Cs: central sulcus. IPs: intraparietal sulcus. syf: sylvian fissure. sts: superior temporal sulcus. Ls: lunate sulcus. IOs: inferior occipital sulcus. Cgs: cingulate sulcus. POs: parietal-occipital sulcus. POm: medial parieto-occipital sulcus. calc: calcarine.

We then examined the brainwide connectivity distributions and their similarity to published anatomical connectivity. Using D99 (version 1.2b)^33^ parcellation, brain areas were classified largely by function into cingulate, insula, orbitofrontal (OFC), lateral prefrontal (Lat. PFC), parietal (Par.), motor (Mot.), auditory (Aud.), somatosensory (Som.), visual occipital (Vis. O.), and visual parietal (Vis. P.), and visual temporal (Vis. T.) (see listing of areas at bottom of **Fig. 3A**). Although the stimulation sites sample only a small portion of the amygdala, overall, the distribution of functional connections (**Fig. 3A**, upper 6 rows in red, rows 1-3 for Monkey K and rows 4-6 for Monkey M) is largely in line with the known distribution of anatomical connections (**Fig. 3A**, lower 6 rows in blue). For example, we found stimulation areas that confirmed prediction based on the know major anatomical connections of the amygdala with the insular areas Ig, Id, and Ia, orbitofrontal and prefrontal areas 11,11, 12, 13 and 14, and visual association areas TPO, TEO, TE, and TAa in the upper bank of the sts. Surprisingly, the connectivity less predicted based on anatomical studies with the exception of premotor area F2, but consistent across the subject monkeys M and K were with motor and premotor areas.

**Fig. 3.**
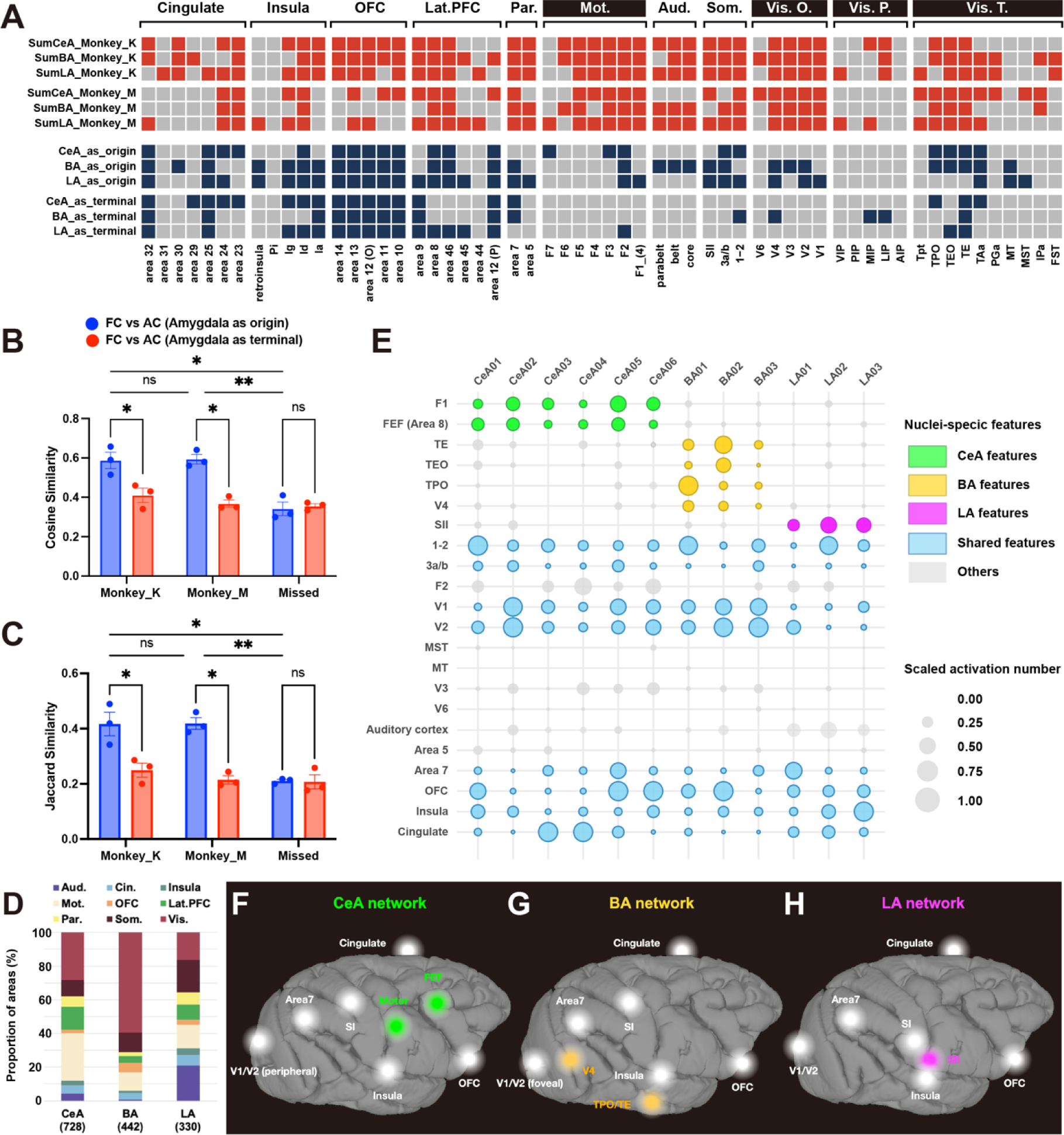
Cortical distributions of CeA, BA and LA networks. (A) A comparison of connectivity revealed by INS-fMRI and by anatomical tracers. Upper 6 rows: red represents presence of functional connections in monkey K and monkey M. Lower 6 rows: blue represents compiled results of anatomical connections originating from the amygdala and anatomical connections to the amygdala originating from the cortex (based on http://cocomac.g-node.org, see method). T.: visual system (temporal). Vis. P.: visual system (parietal). Vis. O.: visual system (occipital). Som.: somatosensory cortex. Lat-PFC: lateral prefrontal cortex. Par.: parietal cortex. OFC: orbital frontal cortex. Mot.: motor cortex. Aud.: auditory cortex. Pi: parainsula. Ig: granular insula. Id: dysgranular insula. Ia: agranular insula. (B-C) Jaccard similarity and cosine similarity between FC (functional connectivity) and AC (anatomical connectivity), the red color represents similarity between FC and AC to the amygdala originating from the cortex, the blue color represents similarity between FC and AC that originating from the amygdala. Results are from 2 monkeys (monkey K and monkey M), and from stimulations sites out of the amygdala (missed) in monkey K. (D) Proportional composition of cortical connections from CeA, BA and LA in monkey K (e.g., for all stimulation sites in CeA, the #voxels in an area connected to CeA / total voxels connected to CeA). (E) Global distribution of activation evoked by different stimulation sites. Each column illustrates activation from a single site (6 in CeA, 3 in BA, 3 in LA). The colors represent that the brain areas have outstanding and consistent activations from stimulating sites in CeA (green), BA (yellow), LA (magenta). Blue color represent that all stimulation sites evoke activations in this brain area. The size of bubbles represents the number of voxels evoked by each stimulation site in each brain area. The numbers were scaled by each stimulation site. (F-H) Summarized global networks involving CeA (F), BA (G), and LA (H), respectively. The colored nodes represent areas dominated by CeA, BA, or LA, the white nodes represent areas receiving similar prominence of connections from CeA, BA, and LA.

To systematically compare the congruence between functional connectivity and anatomical connectivity, we compared the amygdala functional connectivity matrices of two monkeys with anatomical matrices of two directions (axons to/from amygdala), calculating their cosine similarity (cosine of angle between two vectors, **Fig. 3B**) and Jaccard similarity (size of intersection divided by size of union, **Fig. 3C**). Additionally, we computed the functional connectivity matrices for stimulation sites outside the amygdala (missed) as a control. The results demonstrated that the amygdala functional connectivity matrices of both monkeys exhibited greater similarity with the anatomical matrices summarizing axonal connections originating from the amygdala, in contrast to those terminating in the amygdala. We note that this result is consistent with previous observations that stimulation evoked connectivity is biased towards ‘anterograde’ directions^21,34,35^. Thus, the areal connectivity of INS stimulation is consistent with previous anatomical studies based in relatively large injection sites; however, the primary and new finding is the mesoscale patchiness of connectivity.

There were also substantial differences. The matrix reveals that functional connectivity is present in some areas where anatomical connections are not prominently observed, such as the motor cortex, and areas on visual pathways and visual cortex (**Fig. 3A**, marked by white text on a black background), potentially reflecting secondary connections. Another difference lies in the degree of ‘common connectivity’ that CeA, BA, and LA shares with single cortical areas. For example, all three subdivisions exhibit extensive bidirectional connections with the OFC. However, anatomical data reveal limited shared connectivity of CeA, BA, and LA with motor cortex, somatosensory cortex, and visual cortex (**Fig. 3A**). The differences in proportional connectivity reported in anatomical studies and this study are likely due to (1) the secondary connections revealed by INS and (2) the small (mesoscale) size the volume of stimulated projection neurons.

We also compared the relative distributions of connectivity between the three nuclei. Out of the total number evoked from all stimulation sites of each nucleus (CeA: 728, BA: 442, and LA: 330), the percentage of connections associated each different brain area (grouped by function) (**Fig. 3D**). We noted a few distinct characteristics in the three distributions. Relative to BA and LA, CeA had the most functional connections with the motor cortex (28.4%) and the lateral PFC (13.6%). Relative to CeA and LA, BA had the most with the visual cortex (59.6%). And, relative to CeA and BA, LA had the most with the auditory cortex (21.0%) and the somatosensory cortex (19.2%). In contrast, CeA, BA, and LA exhibited similar proportions of functional connectivity with the cingulate cortex (∼5%).

To investigate the nature of the connections in finer detail, we evaluated the relative prominence of activations in different cortical areas from each of the stimulation sites in CeA, BA, and LA (**Fig. 3E**). To compare the global distribution of connections across stimulation sites and to identify the unique or shared features of stimulation sites from different nuclei, we calculated the activation number (# voxels) in each brain area for each stimulation site. To make the data more comparable between sites, we scaled the numbers by stimulation sites using: scaled x = (x – min(x)) / (max(x) – min(x)). max(x) and min(x) are maximun and minimum values for each stimulation site. As seen in the top 9 rows, some areas are dominated by connections from CeA (green: Primary motor cortex [F1], Frontal eye field [FEF]), BA (yellow: TE, TEO, TPO, V4), and LA (magenta: Secondary somatosensory cortex [SII]). Other cortical areas connected with amygdala have significant contributions from all three nuclei (sensory areas 3a/b, 1-2, V1, V2, motor F2, and area 7, OFC, insula, cingulate, represented in blue). Yet other areas have relatively weak connections with amygdala (gray: V3, MT, MST, area 5, V6). These distinctions are summarized on brain views (**Fig. 3F-H**). Thus, the amygdala is extensively connected, directly or indirectly, with most cortical areas; from these sites of stimulation, it appears that dorsal and mediodorsal pathways are more weakly connected; this is consistent with anatomy for area 5 and V6, but not MT, MST, and V3. However, it is possible we did not stimulate the sites connected to these areas.

### Similar but distinct functional connectivity in cingulate cortex, insula, and OFC

Three of the major cortical connections of amygdala include the cingulate cortex, insula, and the orbitofrontal cortex^8^. Here, we illustrate the connections of these sites with the CeA, BA, and LA nuclei of the amygdala (**Fig. 4**). The first general observation is that within each of the activated cortical areas, connected sites appear patchy. For CeA stimulations (**Fig. 4B, 4G, 4L**), patchy activations were observed in three areas: (1) cingulate areas 23, 24, and 32, and medial orbitofrontal area 10, (2) insular areas in the lateral sulcus (lg, ld, lapl) as well more infra-orbital insular areas lai and lal, (3) lateral prefrontal areas 12, 44, 45, 46, as well as (4) infraorbital areas 11 and 13. The stimulation of BA sites (**Fig. 4C, 4H, 4M**) and LA sites (**Fig. 4D, 4I, 4N**) elicited similar activation profiles in the cingulate, insular, and PFC (BA activations in cingulate and insula are consistent with previous study ^28^). Some notable differences include: (1) in contrast to stimulation of CeA and BA, LA shows little activation in area 32 (see also **Fig. 3A**). (2) Stimulation of BA shows little activation in anterior insular area Iai or in posterior Ig. However, closer inspection reveals that the respective activations of CeA, BA, and LA are largely non-overlapping within each area (**Fig. 4E, 4J, 4O**, see merged). Overall, while this spatial comparison shows CeA, BA, and LA shares common cortical targets, the mesoscale connectional architecture comprises largely distinct patchy territories within each cortical area, suggesting a degree of functional segregation.

**Fig. 4.**
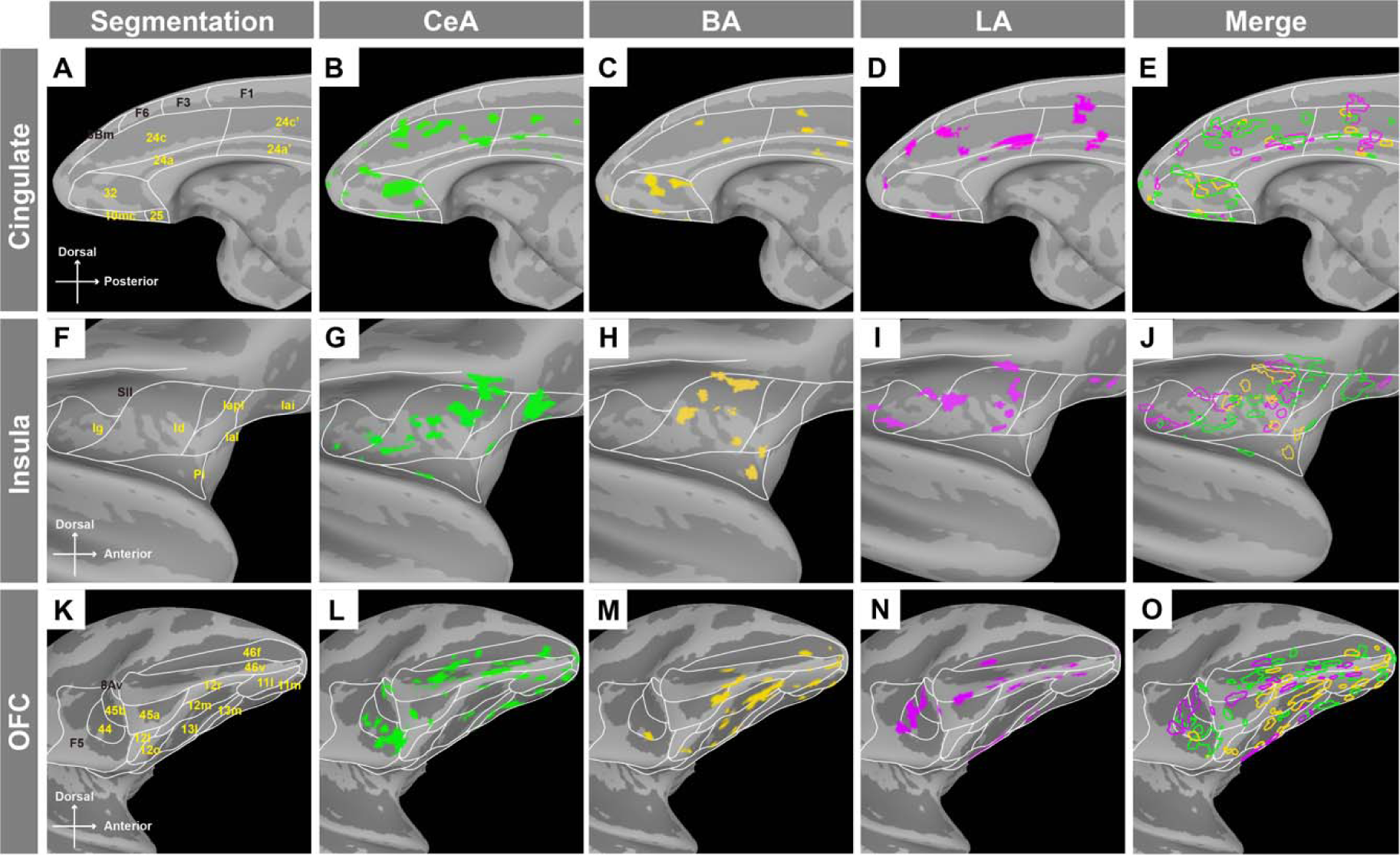
Functional connections with cingulate cortex, insula, and OFC. Topography of connected areas in cingulate cortex (B-E), insula (G-J), and OFC (L-O). Segmentation of the brain areas are shown in the first column (A, F, K). Merged views of CeA, BA and LA are shown in the last column (E, J, O). Iai: intermediate agranular insula. Iapl: posterior lateral agranular insula. Ial: lateral agranular insula. The results are masked by cingulate, insula and OFC for the purpose to highlight results in these areas.

### Mesoscale cortical connection patterns from each of CeA, BA and LA

We next examined the connections from different stimulation sites within each of CeA, BA, and LA to single cortical areas. These regions were selected based on the presence of activation, for a given cortical area, across multiple stimulation sites (as shown in **Fig. 3E**). As shown in **Fig. 5A**, the six stimulation sites in CeA all led to activations in primary motor cortex (**Fig. 5B**) and area 8 of FEF (**Fig. 5C**); these showed distinct and largely non-overlapping distributions that appeared to be topographically organized. The patches corresponding to the four stimulation sites in posterior parts of CeA (CeA01, 02, 03, and 04) were mainly distributed in the lateral part of the primary motor cortex (**Fig. 5B**), potentially corresponding to the head and face motor areas^36,37^. In contrast, the connections corresponding to the two anterior CeA stimulation sites were located more medially in primary motor cortex, possibly in the hand motor area. Interestingly, the activations in primary motor cortex arising from the spatially closer stimulation points (CeA01 & CeA02, CeA03 & CeA04, CeA05 & CeA06, each less than 1mm to each other) were also closer to each other on the cortical surface (**Fig. 5B**). For the FEF (**Fig. 5C**), most connection sites were located within area 8Ad and also exhibited little overlap; however, unlike the motor cortex, there appears to be some interdigitation of the activations from CeA anterior (CeA01 & CeA02) vs. CeA posterior sites (CeA05 & CeA06). In a second monkey (monkey M), patchy activations were observed in both F1 and FEF (**Fig. S4A-C**).

**Fig. 5.**
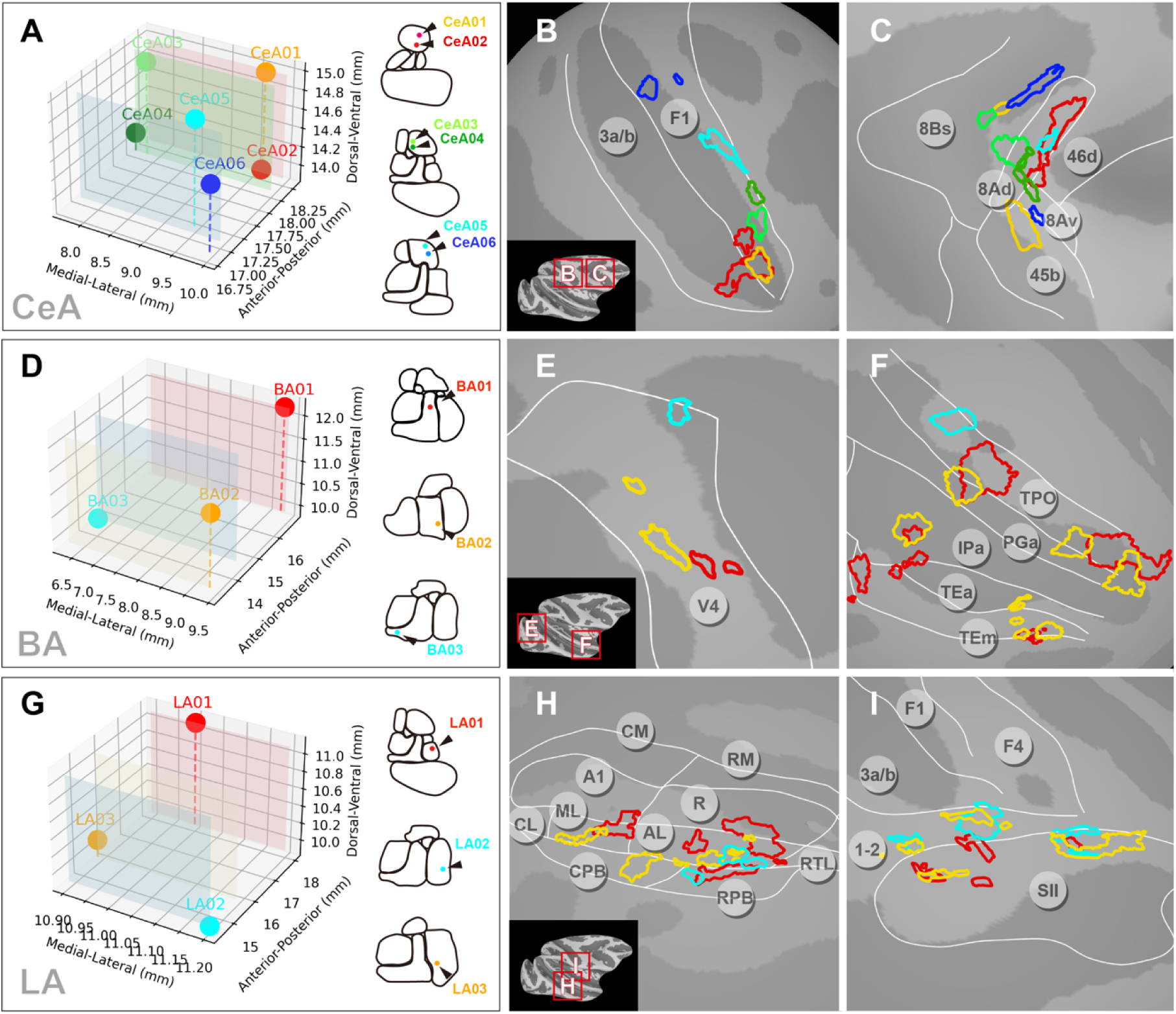
Local cortical topography of connections from single amygdala nuclei. Activations from different stimulation sites within each of CeA, BA, and LA were mapped onto the cortical surface (P<1×10-3). (A, D, G) Stimulation sites are shown in 3d coordinates (left) and in rostro-caudal contour cartoons (right). (A-C) Six stimulation sites in CeA revealed connected sites mostly in F1 (B) and FEF Area 8 (C). (D-F) Three stimulation sites in BA revealed connected sites in area V4 (E) and in ventral visual pathway TP, PG, IP, TE (F). (G-I) Three stimulation sites in LA revealed connected sites in auditory belt/parabelt areas AL, ML, CPB, RPB (H) and somatosensory areas 1-2 and SII (I). A1: primary auditory area. R: rostral area. CM: caudomedial belt region. AL: anterolateral belt region. ML: middle lateral belt region. RPB: rostral parabelt region. CPB: caudal parabelt region.

The connections corresponding to the three stimulation sites in the BA (**Fig. 5D**) led to activation of several patches in V4 with the red patches most lateral (foveal), the cyan patch most medial, and the yellow patches intermediate (**Fig. 5E**), suggesting some foveal to parafoveal topography. In the temporal lobe, area TPO contains alternating red and yellow patches anteriorly, indicating an interleaved pattern of connectivity, as well as a single cyan most posterior patch (**Fig. 5F**). Similar, though smaller, red and yellow patches were observed in TE and IPa. In a second monkey, patchy activations were observed in V4 and TPO/IPa (**Fig. S4D-E**).

The connections from the three stimulation sites in LA (**Fig. 5G**) showed both overlapping and interleaved distributions in higher order sensory areas, including auditory belt and parabelt regions (**Fig. 5H**) and secondary somatosensory cortex (**Fig. 5I**). As shown in **Fig. 5H**, connections in auditory belt cortex were largely distributed in patchy fashion in AL (where red, yellow, and cyan could be viewed as interleaved), with a few additionally distributed in ML, CPB and RPB. For somatosensory cortex (**Fig. 5I**), the connections are mainly distributed along the border of areas 1-2 (yellow and cyan interleaved) and secondary somatosensory cortex, with some overlaps in the connections of the three stimulation sites. In a second monkey, patchy activations were observed in auditory and somatosensory (SII, 1-2) areas (**Fig. S4G-I**).

In sum, we observed that stimulation of different sites within each of CeA, BA, and LA resulted in patchy activations in connected cortical areas. Patches were largely distinct and non-overlapping, and exhibited distinct types of topography (e.g., topographic, interleaved). Data from a second monkey supported the patchy nature of activations. These examples also illustrate that single nuclei within amygdala may have a topographic relationship with one cortical area (e.g. BA with V4) and an interleaved pattern with another (e.g. BA with TPO).

### Mesoscale connection patterns from CeA, BA and LA to single cortical areas

We then examined how CeA, BA, and LA connected to the same cortical area. As shown by the matrix of functional connectivity in **Fig. 3E** (blue bubbles) and **Fig. 3F-H** (white nodes), the brain areas that were strongly activated by all three nuclei include V1/V2 (**Fig. 6A**), SI/SII (**Fig. 6B**), and area 7 (**Fig. 6C**).

**Fig. 6.**
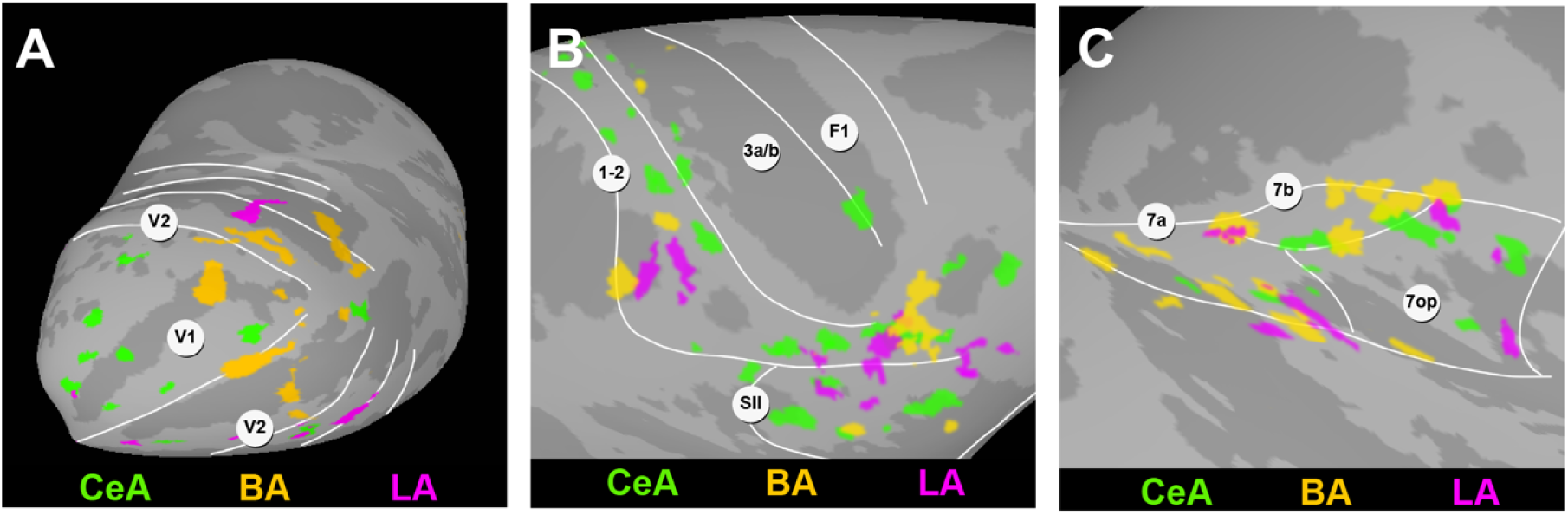
Connectivity patterns in cortical areas with activations from CeA, BA and LA. Topography of connected sites in V1/V2 (A), SI/SII (B), and area7 (C), respectively.

In V1 and V2 (**Fig. 6A**), BA’s connections appeared heavily biased towards the foveal region (on the lateral convexity, yellow) and connections from CeA were predominantly localized in the peripheral regions (green), consistent with an association of BA with foveal visual attention related behavioral circuits and CeA with physiological reaction to peripherally appearing stimuli (see Discussion). Connections from LA are fewer and distributed in a patchy pattern in V1, V2 and V3 (magenta). Notably, the connections from each nucleus are non-overlapping. In SI/SII (**Fig. 6B**), connections with LA were seen at the border between SI and SII and in the facial sensory area of area 1-2 (magenta), while the connections of BA (yellow) were distributed primarily in areas of area 1-2 corresponding to face and anterior parts of SII. Connections with CeA (green) were more broadly distributed across SI and SII. Patches were largely non-overlapping. For area 7 (**Fig. 6C**), the connections from CeA, BA and LA were also largely non-overlapping, with BA more dominant in 7a and 7B, LA in 7a and 7op, and CeA in 7a and 7b.

## Discussion

Our study examines the cortical activations obtained by focal INS stimulation of sites in BA, LA, and CeA nuclei of the macaque amygdala revealed by mapping in 7T fMRI. The matrix and the details of the activation patterns can be summarized in four main points: (1) *Broader networks than anatomical connectivity:* INS-fMRI mapping reveals both primary and secondary functional connections, beyond the direct connections revealed by anatomical tracer studies, thereby providing a brainwide view of functional networks (**Fig. 3A**). In addition, matrix comparison of amygdala-cortical connections revealed by INS and known anatomical connectivity revealed similarities primarily with ‘amygdala as the origin’ (**Fig. 3B-C**), consistent with a the ‘anterograde’ bias of the INS method^21^. (2) *Mesoscale activations:* Activations in cortical connected sites are consistently mesoscale (primarily one to few millimeters) in size (**Fig. 2B**), raising the possibility of functionally specific interfaces with mesoscale cortical architectures. (3) *Functional segregation:* Generally, the mesoscale patches (both those from the same stimulation site and from sites in CeA, BA, and LA) are non-overlapping (**Figs. 3-6**, **Fig. S4**), and can exhibit topographic as well as interleaved arrangements (**Fig. 5-6**), indicating some degree of functional segregation of connections in target cortical areas. (4) *Area-specific integrations:* While our stimulation sites represent only a sample of three nuclei in the amygdala, the current sample shows that some cortical areas interface more strongly with CeA (e.g. F1, FEF), BA (e.g. V4, TE, TEO, TPO), and LA (e.g. SII) (**Fig. 3F-H**), while other cortical areas receive inputs from all three CeA, BA, and LA nuclei (**Fig. 3E**, **Fig. 6**). Cortical areas with strongest connectivity included limbic areas cingulate, insula, and infraorbital cortex (**Fig. 4**), and the somatosensory cortex, visual cortex, and area 7 of the parietal lobe (**Fig. 6**). Interestingly, the spatial distribution of some cortical connectivity patterns revealed the presence of topographic specificity (somatosensory cortex SI and visual cortex V1) or interleaved distributions (area 7).

### Methodological and Statistical Considerations

As this is a relatively new method for studying functional connections in the brain, it is important to evaluate the methods for identifying remote activations. Normally, causal fMRI studies use FDR corrected p values based on GLM analysis to evaluate statistically significant BOLD response; from this, connectivity between stimulated and connected sites are inferred. Depending on the study, BOLD response evoked at connected sites can be more robust with stronger stimulation paradigms and weaker with more focal or more cellularly specific stimulation; thus, across studies a range of corrected FDR values have been used. For example, an estim-fMRI study in monkeys, where activations span several mm in size, used corrected FDR p values <0.00005^38^, while in a human study estim-fMRI of amygdala, cingulate, and prefrontal cortex, these values were p<0.001^39^. In opto-fMRI studies, values of corrected FDR include p<0.000001^40^, p<0.001^41,42^, and p<0.05 (in a cell-type specific study^43^). In a focused ultrasound-fMRI, corrected values of p<0.02, 0.005, and 0.001 were used^44^. Here, INS is a method that stimulates a small (submillimeter) volume of tissue, activating a focal cluster of connected neurons, and results in a relatively small BOLD signal (see **Fig. 1**). Despite this, using statistics that are well within published standards (FDR-corrected p<0.05, range 1.9×10^−3^-1.4×10^−2^), we reveal consistent mesoscale activations and connection patterns in many cortical areas.

These reliability of these activations are further supported by: (1) half trial vs half trial reproducibility^28^ (**Fig. S2A&C**), (2) stability of activations across thresholds, that suggest these mesoscale activation patterns are not an artifact of specific thresholding (**Fig. S3**), and (3) alternating stimulation of two distinct nuclei lead to two distinct sets of reproducible responses (**Fig. S2B**). Furthermore, we observe a similar-sized modularity across brain areas, across stimulation sites (CeA, BA and LA) and across animals. This modular architecture reveals a complementary (non-overlapping) organization of connections (from CeA, BA, and LA). Together, these point to the presence of non-random, non-artifactual, inherent structure in brain connectivity.

### Brainwide mapping using INS-fMRI

We examined the spatial specificity and organization of the networks revealed by INS and high-resolution fMRI (**Fig. 6**). While “column-to-column” cortical connectivity at local scale within visual and somatosensory cortex are known^34,35^, whether brainwide networks are also so organized has not been established. The INS-fMRI method has allowed us to address the question of “what is the remote functional reach of a ‘single’ mesoscale node?” Hu and Roe^35^ showed that stimulation of a single functional domain (column) in V2 elicits a pattern of intra-areal and inter-areal columns that repeats across different functional modalities in V2, demonstrating a canonical (∼10-12 columns) columnar microcircuit that serves distinct feature modalities. However, whether large scale networks are similarly organized has been unexplored. Our fMRI data suggest the presence of analogous minimal mesoscale circuits. Although the amygdala is not organized into columns, local field potentials recorded from multiple nuclei of the primate amygdala reveal the presence of neural clusters within amygdala that differentially process visual, auditory, and tactile stimuli^19^. This type of mesoscale-to-mesoscale specificity at global scale is still a relatively new concept^28^.

This study contributes to a better understanding of the directional and single-or multi-synaptic aspects of connectivity revealed by a single stimulation site. Previous studies have shown that INS-fMRI is biased towards revealing ‘anterograde’ activations; that is, stimulation of a single cortical node leads to activation of middle layers in anatomically known feedforward connections and activation of superficial and deep layers in known feedback connections^21,34^. Consistent with the ‘anterograde’ interpretation, our method reveals a better match to anatomical data with amygdala as the origin than as a recipient of cortical inputs (see matrix in **Fig. 3A**). For example, anatomical methods reveal projections from amygdala to V2, but not from V2 to amygdala^45,46^. Our stimulations also replicate this finding: stimulation of amygdala produces robust V2 activation, but there is an absence of activation in amygdala following stimulation of V2 (data not shown: 23 sites in V2, 20 trials per site, no activated voxels amygdala). This suggests that the connections from CeA to V2 might be disynaptic, possibly through pulvinar or SC or another cortical area such as V1^45,46^.

This demonstrates that the study of brain connectivity can extend to include second synapse connections^47,48^, providing visualization of multiple brainwide functional networks within single animals.

### The mesoscale architecture underlying multimodal processing networks

We find the functional activations induced by INS stimulation are largely non-overlapping. We bear in mind that (1) our functional activations reveal only the foci of strong connectivity and that there may be functionally connected cells whose connectivity do not reach statistical significance with this method and (2) our sample has targeted only a small percentage (∼10%) of the the total CeA, BA, LA volume (total CeA/BA/LA volume ∼100mm^3^ ^49^, ∼1mm^3^/site, 12 sites in Monkey K 12%, 6 sites in Monkey M 6%). Given this caveat, our data suggests that amygdala outputs to cortical areas are received in distinct mesoscale regions and that these regions can be arrayed in different topographic patterns. These patchy connections bring into focus distinctions that may not be apparent from traditional anatomical tracer injection studies^50,51^, but have been long indicated by anterograde single axon tracing studies^52–55^. This functional stimulation approach offers a way to study and compare the distribution of multiple connectivity sources. While we do not yet understand the significance of the distinct connectional modes (topographic vs interleaved), parallel studies in the visual system may be informative.

For areas that appear to receive inputs predominantly from one of the three amygdala nuclei (Fig. 5A), it could suggest the influence of a single functional domain within the amygdala on the native mesoscale architecture of the recipient cortical area. Although the amygdala has a dozen functional nuclei, autonomic and homeostatic control is strongly associated with the CeA. Growing evidence indicates the role of the amygdala in facial recognition and in the valence and meaning of facial emotions^19,56,57^. CeA mediated coupling of certain motor behaviors, such as eye movements with autonomic changes in response to alerting signals could underlie the association of CeA with F1, FEF, and peripheral visual cortex (**Fig. 3F, 5B, 6A**) (e.g., facial expressions and other social signals through gestures, postures, etc.) (**Fig. 5B**), whereas the association of BA with foveal regions of visual cortex and face regions of areas 1-2 (**Fig. 5E&F, 6A&B**) may be associated with the ventral pathway evaluation of facial gestures. Likewise, the connections of the BA and LA that project in an interdigitating fashion to temporal areas, may impact distinct object-based (e.g., face patches) or sensory-based modalities in ventral visual pathway (TPO/TE/V4) and sensory cortex (auditory belt areas, somatosensory areas 1-2/SII), respectively (**Fig. 5E, F, H**). The critical role of these amygdalofugal projections for the functionality of the temporal cortical areas has been demonstrated by comparing the activation of multiple face-responsive visual areas in temporal cortical areas before and after excitotoxic lesions of the amygdala. In the absence of inputs from the amygdala these cortical areas failed to respond to the stimuli that activated them reliably before the lesion^58^.

The connections appear to be segregated in areas that receive inputs from multiple nuclei, such as early visual cortex, central visual fields may be dominated by BA, while extrafoveal regions by CeA influences (**Fig. 6A**) or by integration of the amygdala inputs in face vs. body areas (**Fig. 6B**). Areas such as parietal area 7 exhibit a highly interdigitating pattern of CeA, BA, and LA inputs, potentially indicating a high degree of limbic, cognitive, and sensory integration for modulating spatial transformations of behavioral maps (**Fig. 6C**). These several patterns of connectivity suggest that amygdala’s influence on brainwide networks is mediated in an organized functionally specific manner.

These findings align with two features of our current understanding of cortical function: 1) Some cortical function rely on spatial topography such as cortical columns in the visual cortex^59^ or stripes distributed across the motor cortex^60^. There is also evidence that within motor cortex the topography for motor action interdigitates with regions for combining action and physiological functions such as arousal and pain^37^. Neurons aggregated in the same cortical functional domain share a functional processing goal (e.g., color, shape, disparity, motion in visual cortex, action vs interoceptive nodes in motor cortex, object vs face patches in temporal cortex). Thus, the connections of the amygdala to different units may indicate the amygdala’s processing of various features through different internal neuronal clusters. 2) There is integration of multiple sensory and motor functions. Such integration might occur within the functional cortical areas, or in higher-order cortices, or in subcortical structures. Each stimulation site we targeted is connected to multiple cortical areas representing the different axes of behavior, indicating the amygdala plays a significant bridging and integrative role in the emotion-cognitive-sensation-motor circuit.

Taking a step further, we suggest that the mesoscale networks comprise a scaffold upon which dynamic modulation is conducted. That is, within each node are related neurons which share common targets. For example, viral barcode analysis of amygdala-prefrontal connectivity has shown that single amygdala BLA neurons can connect with one to several neurons in different parts of prefrontal cortex^61^. This raises a potential scenario in which dynamics of mesoscale nodal selection is further coupled with intra-node single neuron selection to achieve a broad range of distinct and specific functional circuits. In this manner, highly organized and sparse mesoscale networks may still achieve a rich repertoire of integrative yet specific affective behaviors. Further studies using different intensities of stimulation in controlled behavioral contexts may test this proposal.

### Comparison with previous studies

To understand the connectivity patterns of the amygdala, the earliest and most direct method employed neural tracers to study connections from an anatomical perspective. This approach led to the recognition of the prominent connections between the amygdala and OFC, insula, and anterior cingulate cortex, and many other cortical and subcortical areas, establishing the structural basis for the affective modulation of multiple functions. Building on this foundation, studies employing electrical stimulation and neurochemical modulation have provided further understanding of effective connectivity of the amygdala; these revealed a broader set of functional connections, including those with the posterior cingulate, retrosplenial cortex, parietal cortex, and temporal cortex^16,62^. At the whole-brain level, neurochemical modulation of the amygdala via designer receptors exclusively activated by designer drugs (DREADDs) has shown significant impacts on global brain networks^62,63^. Research conducted on stereotactic electro-encephalography in epilepsy patients, through direct electrical stimulation, has observed connected areas shared by BA and LA, including OFC, insula, anterior cingulate cortex, and post-central gyrus, revealing temporal and spatial differences in the connectivity patterns of different nuclei^16^. Another study^64^ applied electrical stimulation in awake epilepsy patients and evaluated the patients’ sensations, revealing the integrative role of various nuclei in mediating emotional reactions and sensory functions including visual, auditory, and vestibular sensations. These studies indicated that the modulation of the amygdala affects not only areas directly connected to it but also the activity of secondary regions. Similar to existing anatomical findings, we observed connections between the amygdala and the OFC, insula, and cingulate gyrus; in addition, focal connections with multiple areas, including the somatosensory, auditory, visual, and motor cortices, exhibited distinct topographic mesoscale organizations. These topographies appeared to fall broadly into three classes described as parallel, interdigitating, and convergent. Our study thus echoes and extends previous findings, revealing the fine-scale organization of how different axes of amygdala function (BA, LA, CeA) influence individual cortical areas as well as selectively integrate brainwide circuits for emotion-guided social behavior.

## Materials and Methods

Methods used here for macaque monkey animal procedures, amygdala INS stimulation, data acquisition and analysis are similar to that described in^28^.

### Macaque monkeys

Two hemispheres in two Rhesus macaques (Macaca mulatta) were used (Monkey K: right amygdala, Monkey M: left amygdala). We have analyzed and present here 12 stimulation sites from 12 sessions from Monkey K (see **Fig. 2A**), and 6 stimulation sites from 6 sessions in Monkey M (see **Fig. S1**, **Fig. S4**).

### Animal preparation and surgery

All procedures were in accordance with the National Institute of Health’s Guide for the Care and Use of Laboratory Animal and with the approval of Zhejiang University Institutional Animal Care Committee. In an initial session, high resolution structural and vascular scans were obtained. Sites to be targeted in the amygdala were then planned and, a grid was implanted in over one hemisphere to aid in the systematic targeting of multiple sites in different nuclei of the amygdala. The animals were sedated with ketamine hydrochloride (10 mg/kg)/atropine (0.03 mg/kg) and anesthetized with 1-2% isoflurane; then, the animals were intubated, placed in a custom MR-compatible stereotaxic apparatus, and artificially ventilated. After local infiltration of skin with lidocaine 1%, a small incision was made in the scalp and a small burr hole craniotomy was then performed at one of the grid site locations determined by previous structural scans for targeting CeA, BA, and LA. During the entire procedure the animal’s body temperature was maintained at 37.5-38.5 °C with a water blanket. Vital signs (heart rate, SpO2, end-tidal CO2, and respiration rate) were continuously monitored. During the scan, monkeys were maintained with sufentanil (2 to 4 μg/kg per hour CRI (continuous rate infusion); induction, 3 μg/kg supplemented with 0.2-0.5% isoflurane). Vital signs (heart rate, SpO2, end-tidal CO2, respiration rate, temperature) were continuously monitored. Following data acquisition, the chamber was cleaned and closed, and animals recovered. Single sessions were conducted once every 1-3 weeks. For terminal experiments in monkey K (which lasted 2-3 days), following completion of data collection, the animal was given an overdose of euthanasia agent iv.

### INS stimulation paradigm

To determine the position of the tip within the amygdala, we conducted a structural scan prior to every INS stimulation run, which revealed a dark spot of signal dropout distinct from surrounding tissues (see **Fig. 1C**). Stimulation sites were further confirmed by location of fiber tip BOLD activation. We applied INS paradigms previously shown to be effective at neuronal activation. As in our previous studies^21,28,29^, INS stimulation (see **Fig. 1**), each trial consisted of 4 pulse trains (12 sec) followed by 18 sec to allow the BOLD signal to return to baseline. Each pulse train lasted 0.5 sec (100 pulses, pulse width 250µs, delivered at 200Hz), with 2.5 sec between each of the 3 pulse trains. This quadruple of pulse trains was delivered once every 30 seconds and repeated 15 times (15 trials) for each run, 1 intensity per run (total period of 450 sec). Radiant exposures which were previously shown to be non-damaging^26,27^ ranged from 0.1-0.5 J/cm2. For most of the runs, we used the stimulation intensity of 0.2 J/cm2. The stimulation intensity was consistent during each run. Typically, 2 runs were conducted per site using 0.2 J/cm2 intensity.

### Data acquisition procedure

Functional images of voxel size 1.5-mm-isotropic were acquired in a 7-Tesla Magnetom MR scanner (Siemens, Erlangen, Germany) with a customized 6-channel receive coil (inner diameter 6-7 cm) with a single-loop transmit coil (inner diameter 18 cm) and a single-shot echo-planar imaging (EPI) sequence (TE 25 ms; TR 2000 ms; matrix size 86 × 72; flip angle 90°). This coil provided improved homogeneity of temporal signal-to-noise ratio (tSNR) over regular surface coils, resulting in images with similar tSNR values (mean tSNR of gray matter ∼75). Functional images from opposite phase-encoding direction were also acquired for correction of image distortion^65^. In addition, Magnetization Prepared Rapid Acquisition Gradient-Echo (MPRAGE) sequence was used to get structural images of voxels size 0.3 mm (monkey K) or 0.5mm (monkey M) isotropic.

### Detection of significant responses

Structural and functional images in raw DICOM files from Siemens scanner were converted to NIfTI^66^ and AFNI (Analysis of Functional NeuroImages) format^67^. Functional images were preprocessed with correction for slice timing, motion, image distortion and baseline shift. Significant responses were identified in a commonly used generalized linear model (GLM) approach, in which the timecourse of each voxel was regressed on the stimulus predictor (see Fig. 1). The stimulus predictor was the convolution between laser onsets and the standard hemodynamic response function. Regression coefficients were subjected to T-tests. The BOLD signals presented in **Fig. 1 E&J** were bandpassed at 0.01-0.08 using 3dBandpass. Only voxels with voxels with significant T-test p-values were highlighted on top of the structural images (p<1×10^−3^), the median FDR-corrected (Benjamini-Hochberg) p was 6.1×10^−3^ [range 1.9×10^−3^-1.4×10^−2^]. Individual voxel timecourses were extracted from EPI data with AFNI 3dmaskdump. Timecourses of percentage signal change were calculated at each timepoint tn as: (Signal(tn)-Signal(t0)/Signal(t0). Timecourses were then averaged over repetitions (15 trials) and plotted. Each baseline was estimated with the mean MR signal over full timecourse. The analyses were done with software AFNI (version 21.0.20) ^67^, FreeSurfer (v6.0.0)^68^, Nipype^69^, Bash, R (4.0.2) and Python (3.11.6). Out of all the voxels in the brain, only those voxels with statistically significant correlation with the stimulation site are considered. Significant voxels were visualized on skull-stripped structural images using FreeSurfer v6.0.0 software package (https://surfer.nmr.mgh.harvard.edu/).

### Tests for Reliability

To examine whether these significant sites represent reliable functional connections, several analyses were conducted to support the reliability of the activations. (1) Half and half analysis: To examine which voxels were reliable, runs were divided into two groups (e.g. even and odd runs) and GLM correlation analyses as described above conducted. Similarity of the activation pattern supported the reliability of response (see **Fig. S2**). (2) Alternative stimulation paradigm: To assess the resolution capability of INS for differentiating connected sites in response to varied stimulation sites, we inserted two optical fibers in the amygdala of monkey M, alternately stimulating sites within BA and LA, and conducted GLM analyses on BA/LA trials separately. The results indicated that trials involving stimulation of BA specifically activated TPO, while those stimulating LA specifically activated the auditory cortex (see **Fig. S2B**), mirroring findings from continuous stimulation of either BA or LA sites with a single optical fiber. This reflects INS’s accuracy for spatial investigation of whole-brain networks. (3) Stability across thresholds: Activation maps were examined using different p values (resulting in larger activation sizes with less significant p values). The corresponding activation patterns remain generally stable, reinforcing the reliability of the method functional connectivity between the amygdala and voxels with significant correlation (see **Fig. S3**, Yao et al 2023). (4) Similarity across animals: We compared, across animals, activation patterns following stimulation of the same (or very similar) sites in the amygdala (see **Fig. 5** and **Fig. S4**, Shi et al 2021).

### Image alignment

All structural and functional images were co-registered to the digital version of rhesus monkey atlas with AFNI command 3dAllineate and 3dNwarpApply. We used D99 digital atlas (version 1.2b)^33^ for cortical segmentation, and SARM digital atlas^70^ for subcortical segmentation. The alignment was then manually examined according to an MR-histology atlas^71^, as well as www.brainmaps.org for subcortical and brainstem sites, annotations of brain regions were then assigned to all voxels in the brain. Stimulation sites were determined in structural images on which the tip of the optic fiber was dark and distinct from tissues (see **Fig. 1C**) and in functional images based on functional activation (see **Fig. 1D**).

### Determining voxel number and cortical patch size

We counted the number of voxels in the whole brain (including cortex, subcortical, and brainstem areas), determined using a brain mask (automated, then manually reconfirmed), and then determined by AFNI command 3dBrickStat. The number of voxels activated from each stimulation site, at specific thresholds (p<1×10^−3^), were then determined and percentage out of total voxels calculated. For area-specific voxel counting, we applied the aligned atlas to acquire the certain number of activated voxels in different areas. For calculation of cortical patches, the voxels above the thereshold were first transformed using FreeSurfer command mri_vol2surf and mri_cor2label, and then measured using mris_anatomical_stats.

### Anatomical connectivity matrix

To obtain anatomical evidence of connections between the amygdala and various brain regions, we utilized the CoCoMac database (http://cocomac.g-node.org) to identify axonal projections originating from or terminating in CeA, BA, and LA of the amygdala. Initially, we retrieved comprehensive lists of synonymous text IDs for CeA (62), BA (120), and LA (61). Our inclusion criteria were restricted to sites located within single nuclei, while we excluded sites encompassing multiple nuclei (for example, ‘basolateral’ sites were excluded because they involve both BA and LA). Next, we generate lists of axonal projections by setting amygdala sites as the axon origin sites and axon terminal sites, respectively. Finally, we filtered projections that partially or completely overlapped with the targeted area, supporting the existence of such anatomical connections. For comparison with functional connections (**Fig. 3A-C**), these anatomical connections were manually attributed to corresponding brain regions of the D99 atlas. To compare the similarity between functional connectivity and anatomical connectivity, cosine similarity and Jaccard similarity were calculated between FC (pooled for CeA, BA and LA, respectively) and AC matrix.

### Data visualization

Prism version 8.4.3 for mac, GraphPad Software, La Jolla California USA was used for statistical analysis. Python version 3.11.5, package “matplotlib”^72^, R version 4.0.2, package “ggplot2”, “ComplexHeatmap” were used for data visualization.

## Funding

STI 2030—Major Projects 2021ZD0200401 (AWR)

the National Natural Science Foundation of China U20A20221(AWR)

the National Natural Science Foundation of China 819611280292 (AWR)

the Key Research and Development Program of Zhejiang Province 2020C03004 (AWR)

MOE Frontier Science Center for Brain Science & Brain-Machine Integration (Zhejiang University) (AWR)

the Fundamental Research Funds for the Central Universities (AWR) NIH R01MH121706 (KMG)

## Author contributions

An Ping: Performed surgeries, conducted data analysis, made figures, wrote paper, conducted scans, developed data acquisition and analysis methodology.

Jianbao Wang: Conducted scans, developed methodology, developed data acquisition and analysis methodology.

Miguel Ángel García-Cabezas: Wrote paper, review and editing paper.

Lihui Li: Conducted scans, developed data acquisition and analysis methodology.

Jianmin Zhang: Developed data acquisition methodology.

Junming Zhu: Developed data acquisition methodology, supervise project.

Anna Wang Roe: Obtain funding, design experiments, supervise project and analysis, made figures, and wrote paper.

Katalin Gothard: Obtain funding, made figures, wrote paper, review and editing paper.

## Competing interests

Authors declare that they have no competing interests.

## Data and materials availability

The data to evaluate the conclusions of this study are available within the article and the supplementary materials. Additional data are available on request.

## Figures and Tables

**Fig. S1.**
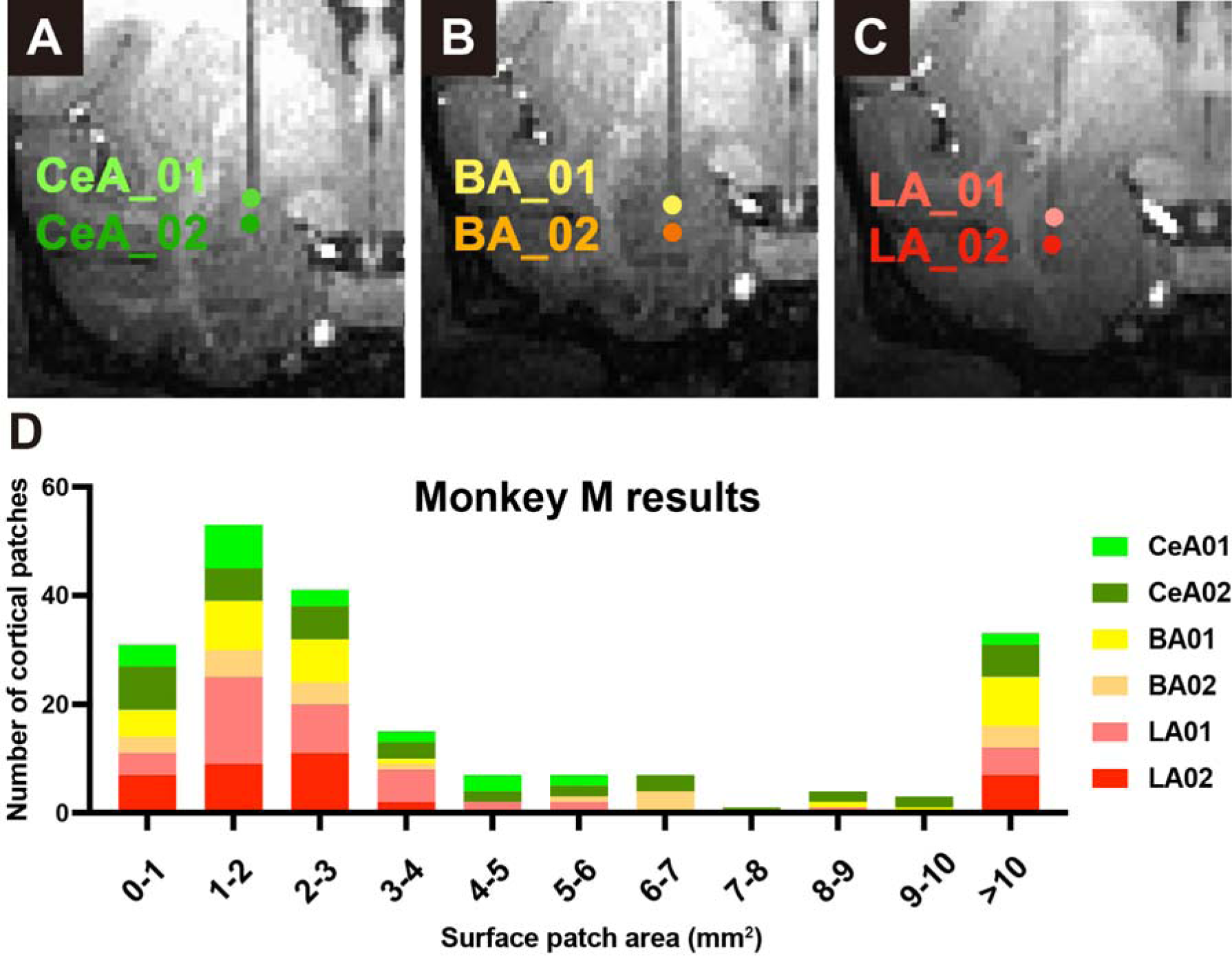
The stacked histogram for patch size of brainwide cortical activations (Monkey M). The x axis represents the size of patches in millimeter square. The y axis represents the number of patches of different sizes. Each color represents the statistics of a stimulation site in monkey M, namely 2 sites in CeA (upper row, shades of green), 2 sites in BA (middle row, shades of yellow) and 2 sites in LA (lower row, shades of red).

**Fig. S2.**
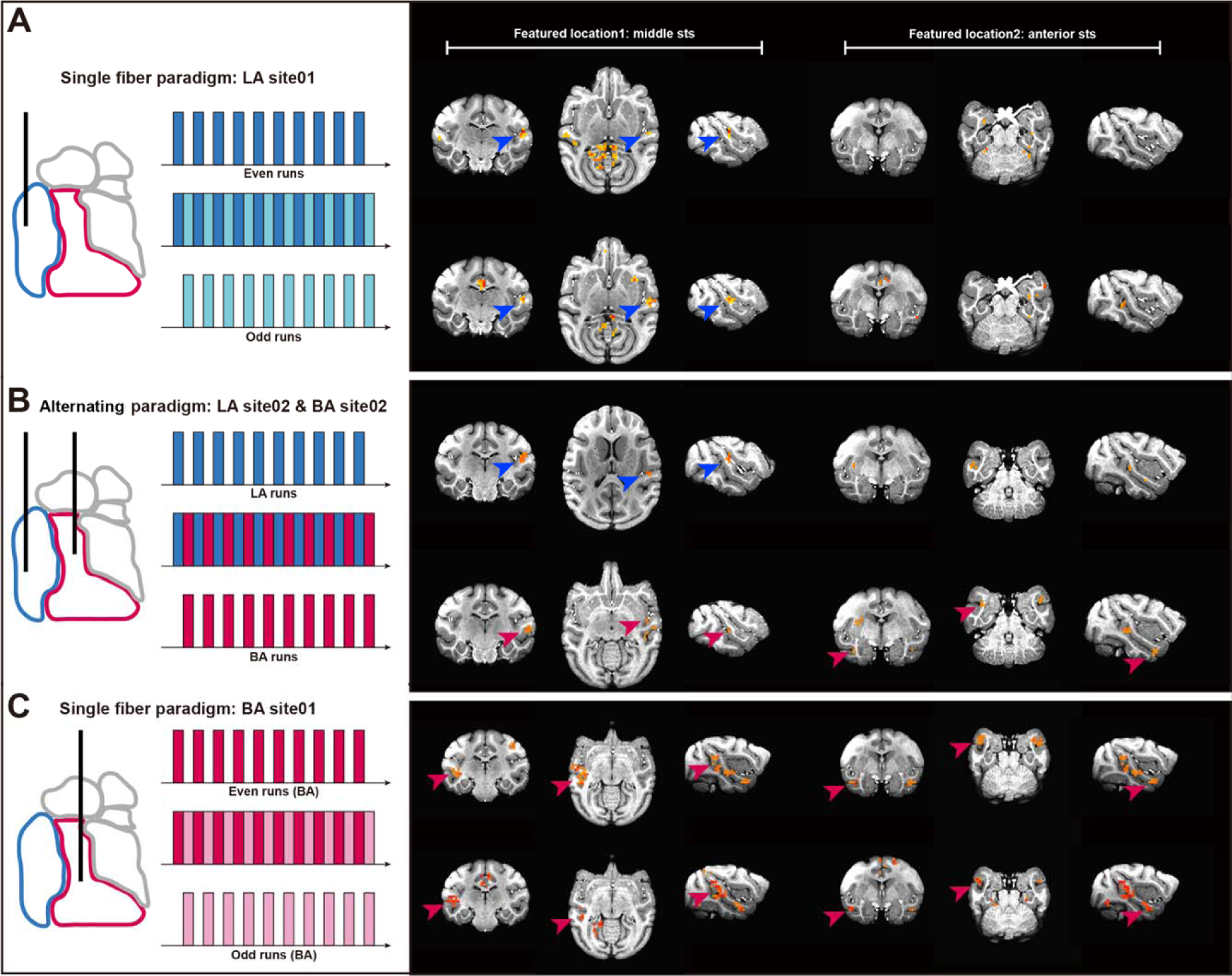
Half and half analysis and alternative stimulation paradigm. We examined the reliability of brainwide activations to amygdala INS stimulation by comparing half trials. Left: half-half analysis (A&C) and alternating stimulation (B) were used. Right: Activations in two regions of the brain are shown (left: middle sts, right: anterior sts). (A) For an example stimulation site in LA, 20 trials were divided into even runs and odd runs and then analyzed using GLM model separately. (B) For a pair of stimulation sites, one in LA and another in BA, the stimulation of each was performed alternatively, each for 10 trials. (C) Same as in (A), except the stimulation site was in BA. The same threshold level was selected for all tests (p<5×10-3). Data from monkey M. The results in (B) indicated that trials involving stimulation of BA specifically activated TPO, while those stimulating LA specifically activated the auditory cortex, mirroring findings from continuous stimulation of either BA (A) or LA (C) sites with a single optical fiber.

**Fig. S3.**
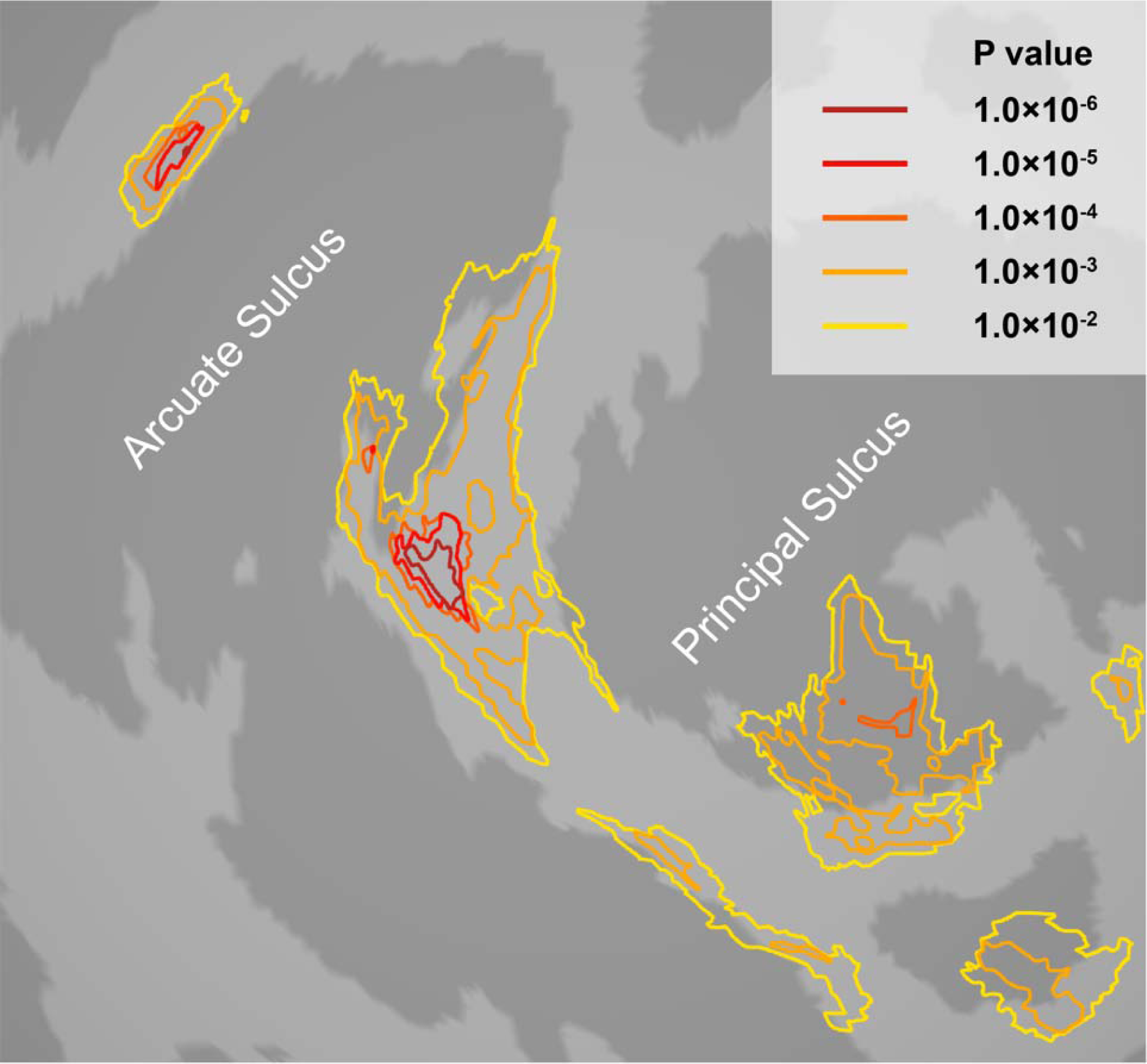
An example of the response patterns at different thresholding p values. The activation evoked by stimulating at a site in medial CeA. The colors of the contours stand for the thresholding percentage. The relationship of thresholds and the corresponding p values are presented at the right upper corner. The main point is that, while the sizes of activations increase with lower threshold, the locations of activations remain largely stable. Our data emphasize the most significant activations seen (highest correlation values), reflecting the ‘backbone’ of the functional network.

**Fig. S4.**
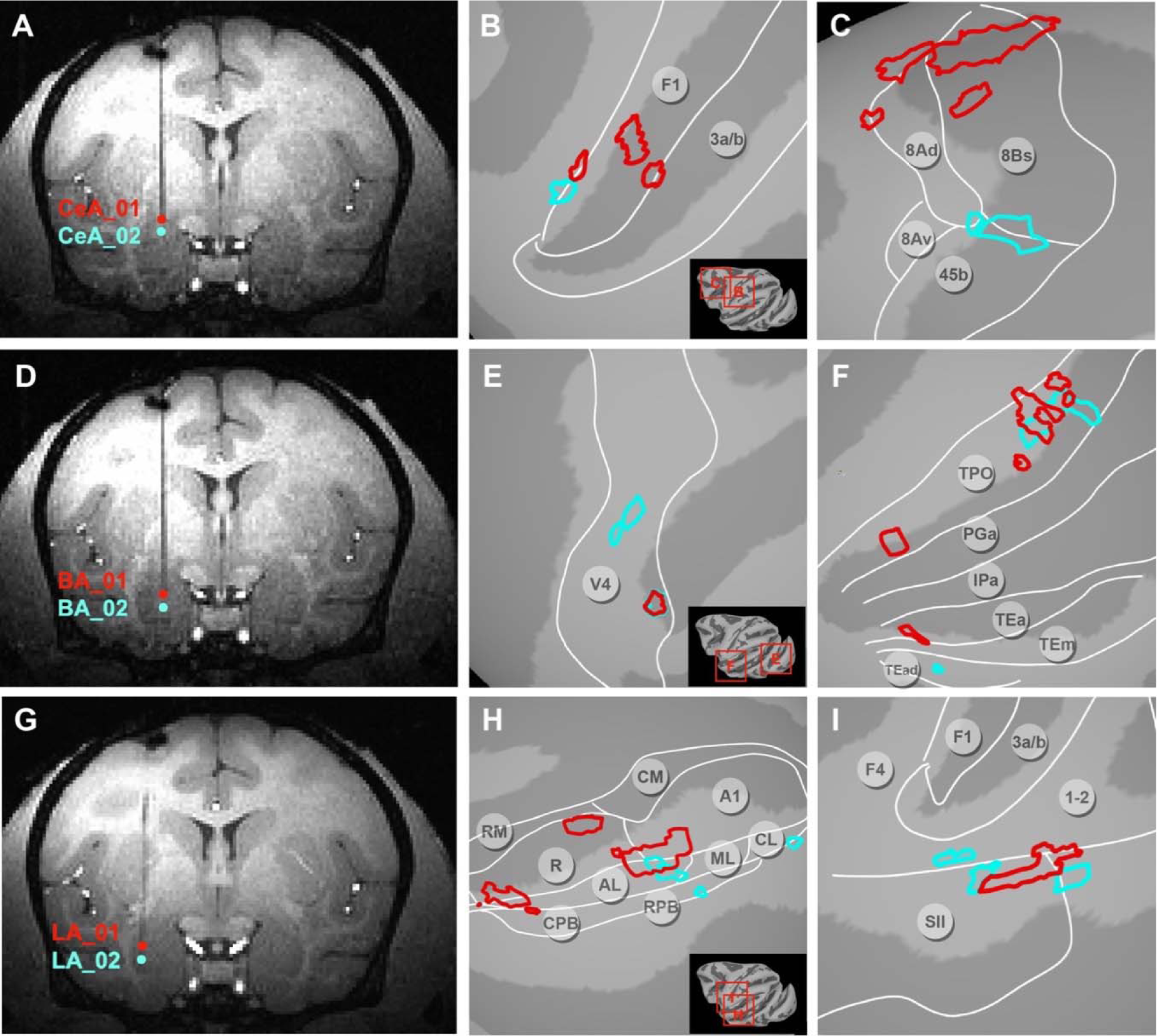
Local cortical topography of connections from the amygdala (Monkey M). (A-C) Two stimulation sites (A) in CeA revealed connected sites in F1 (B) and FEF (C). (D-F) Two stimulation sites (D) in BA revealed connected sites in area V4 (E) and in ventral visual pathway TP, IP (F). (G-I) Two stimulation sites (G) in LA revealed connected sites in auditory areas (H) and somatosensory areas SII (I). Note that the colored patches (P<1×10-3) indicate activation locations and do not contain correlation strength information. Monkey M dataset: right brain.

## Notes

### Competing Interest Statement

The authors have declared no competing interest.

### Summary of Updates

New figure2 and its section updated to provide a generalized result to describe the pattern of mesoscale brainwide connections. Improved figure3 and its section updated for more statistical evidence about the similarity and difference between anatomical and functional connections. Figure4-5 revised. Old schematic figure7 deleted. Discussion updated. Supplemental files updated. Authorship order updated.

